# Structure of the mouse TRPC4 ion channel

**DOI:** 10.1101/282715

**Authors:** Jingjing Duan, Jian Li, Bo Zeng, Gui-Lan Chen, Xiaogang Peng, Yixing Zhang, Jianbin Wang, David E. Clapham, Zongli Li, Jin Zhang

**Affiliations:** School of Basic Medical Sciences, Nanchang University, Nanchang, Jiangxi, 330031, China; Howard Hughes Medical Institute, Janelia Research Campus, Ashburn, VA 20147, USA; Department of Molecular and Cellular Biochemistry, University of Kentucky, Lexington, KY 40536, USA; Key Laboratory of Medical Electrophysiology, Ministry of Education, and Institute of Cardiovascular Research, Southwest Medical University, Luzhou, Sichuan, 646000, China; The Key Laboratory of Molecular Medicine, the Second Affiliated Hospital of Nanchang University, Nanchang 330006, China; Howard Hughes Medical Institute, Department of Biological Chemistry and Molecular Pharmacology, Harvard Medical School, Boston, MA 02115, USA

## Abstract

Members of the transient receptor potential (TRP) ion channels conduct cations into cells. They mediate functions ranging from neuronally-mediated hot and cold sensation to intracellular organellar and primary ciliary signaling. Structures belonging to the TRPV, TRPM, TRPP, TRPA and TRPML subfamilies have been solved, but to date, none of the founding canonical (TRPC) structures. Here we report an electron cryo-microscopy (cryo-EM) structure of TRPC4 in its apo state to an overall resolution of 3.3 Å. The structure reveals an unusually complex architecture with a long pore loop stabilized by a disulfide bond. Beyond the shared tetrameric six-transmembrane fold, the TRPC4 structure deviates from other TRP channels with a unique cytosolic domain, this unique cytosolic N-terminal domain forms extensive aromatic contacts with the TRP and the C-terminal domains. The comparison of our structure with other known TRP structures provides molecular insights into TRPC4 ion selectivity and extends our knowledge of the diversity and evolution of the TRP channels.

## Introduction

Mammalian transient receptor potential (TRP) channels are activated by a wide spectrum of signals -- ligands, temperature, lipids, pH -- and as yet unknown stimuli. They are classified into six subfamilies based on sequence similarity: TRPC (“canonical”), TRPM (“melastatin”), TRPV (“vanilloid”), TRPA (“ankyrin”), TRPML (“mucolipin”), and TRPP (or PKD) (“polycystin”) (1). The TRPC subfamily are non-selective cation channels (Na^+^, K^+^, Ca^2+^) that alter proliferation, vascular tone, and synaptic plasticity (2, 3). This family can be further subdivided into two subgroups: TRPC2/3/6/7 and TRPC1/4/5. TRPC4 is broadly expressed in human tissues and can assemble as homomeric channels or form heteromeric channels with TRPC1 and TRPC5 (4-7). Studies of *Trpc4*-deficient mice have shown that TRPC4 affects endothelial-dependent regulation of vascular tone, endothelial permeability, and neurotransmitter release from thalamic interneurons (8). Stimulation of G_q_ and G_i/o_-G protein coupled receptors (GPCRs) as well as tyrosine kinase receptors potentiate channel activity (9, 10). Activation is regulated by intracellular Ca^2+^, phospholipase C, and membrane lipids by unclear mechanisms.

Along with the revolution in cryo-EM, improved sample preparation, data acquisition, and image processing strategies, the structures of TRPV1 (11-15), TRPA1 (16), TRPP1(17), TRPML1(18) and TRPM4 (19-21) have been solved, but to date, no structures of the canonical (TRPC) family have been published. Here we present the structure of mouse TRPC4 in its apo state at pH 7.5 at an overall resolution of 3.3 Å.

### Overall structure of the mouse TRPC4 tetrameric ion channel

The mouse TRPC4 (residues a.a. 1-758, excluding a.a. 759-974) was expressed using the BacMam expression system (Methods) and purified protein (pH 7.5) was used for single-particle cryo-EM analysis (**Extended Data Fig. 1a).** The conduction properties of TRPC4 currents were identical between truncated and full-length constructs, suggesting that our truncated construct permeates cations (**Extended Data Fig. 1c**). Ca^2+^ measurements and electrophysiological studies were performed to verify that the truncated construct retained sensitivity to channel activators and blockers, as well as responding to GPCR stimulation (**Extended Data Fig. 1b, c**). The overall resolution of TRPC4 reconstruction was 3.3 Å (**Extended Data Fig. 2, Table S1**), which enabled us to construct a near-atomic model (**Extended Data Fig. 3**). Disordered regions led to poor densities for 4 residues in the S1-S2 loop, 2 residues in the S3-S4 loop, 27 residues in the distal N terminus, and 28 residues in the truncated distal C terminus. In total, the TRPC4 structure is a four-fold symmetric homotetramer (**Fig. 1a**) with dimensions of 100 Å × 100 Å × 120 Å (**Fig. 1b**). Each monomer consists of a transmembrane domain (TMD) and a compact cytosolic domain. The cytosolic domain is composed of two subdomains; the N-terminal subdomain consisting of four ankyrin repeats (AR1-AR4) and seven α-helices (H1-H7), and the C-terminal subdomain containing a connecting helix and a coiled-coil domain (**Fig. 1c, d**).

**Figure 1:**
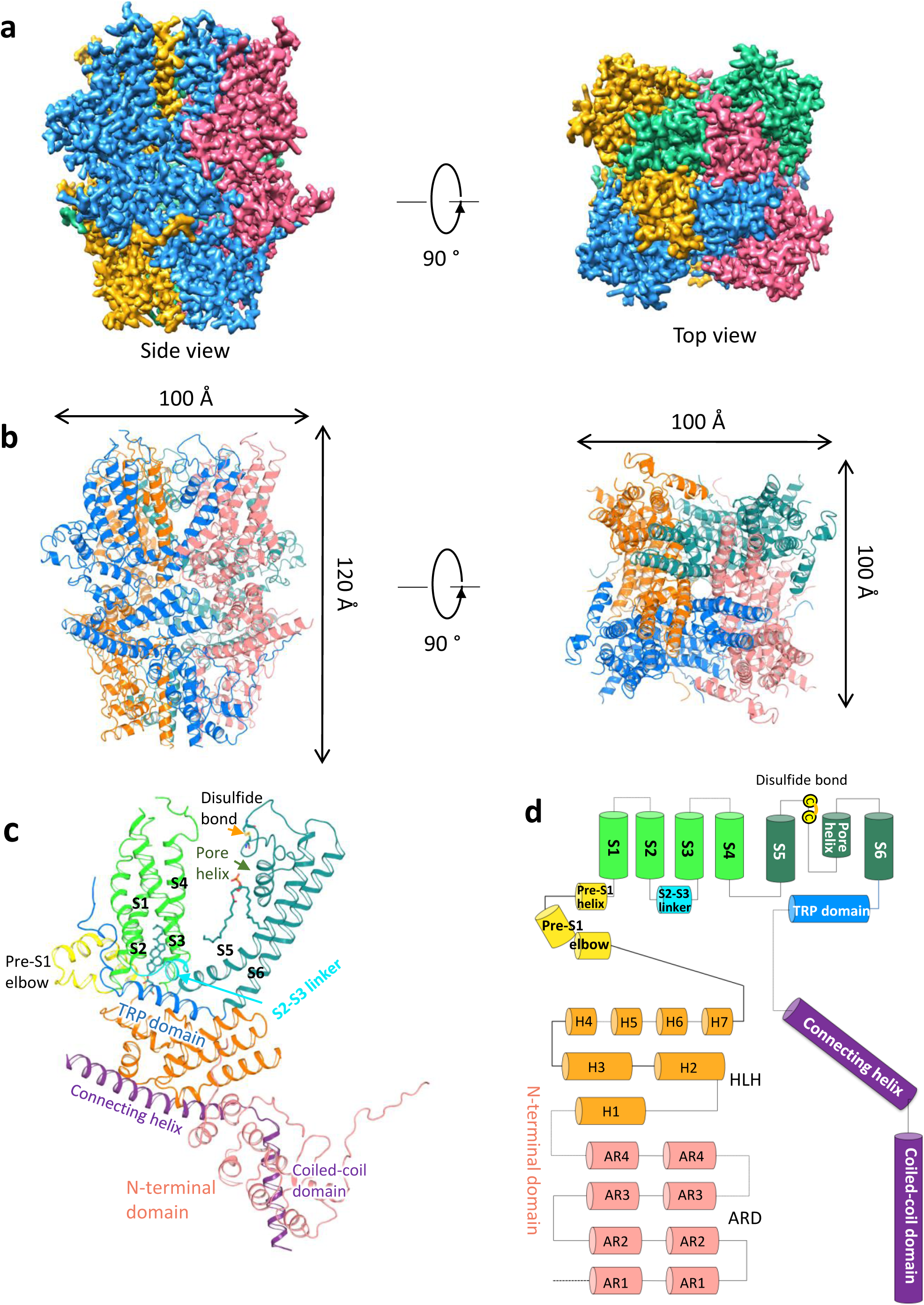
Overall structure of mTRPC4. **a**, Side and top views of cryo-EM density map of mouse TRPC4 at 3.3 Å overall resolution. Each monomer is represented in different colors. **b**, Ribbon diagrams of the mouse TRPC4 model with channel dimensions indicated. **c**, Ribbon diagrams depicting structural details of a single subunit. **d**, Linear diagram depicting the major structural domains of the TRPC4 monomer, color-coded to match the ribbon diagram in c.

### Major structural differences with other TRP subfamilies

In **Fig. 2**, we compare the TRPC4 structure with previously reported TRP structures. Not surprisingly, the organization of 6 helices in each TMD is similar to that of other TRP channels, while the intracellular architecture is distinct. By superimposing a TRPC4 monomer with representative TRP monomers from each subfamily, we found that the overall fold of TRPC4 is closest to that of TRPM4 (**Fig. 2**). TRPC4 has marked similarities to TRPM4 in the TMDs despite their different tissue functions and lack of sequence conservation (<20% identical residues) (**Extended Data Fig. 4**). Distinctive features of TRPC4 include: 1) the arrangement of S2-S3 linker, S5, S6, and the pore loop. In TRPC4, the S2-S3 linker is a two-helical turn, shorter than that of TRPM4 (**Extended Data Fig. 4**), which limits the interactions of S2 and S3 with their cytoplasmic regions; 2) the disulfide bond between TRPC4’s Cys549 and Cys554 lies in the loop linking S5 and the pore helix (**Fig. 2b, c**), while TRPM4’s disulfide bond is located in the loop between the pore helix and S6. Note that these two cysteines are conserved in TPRC1/4/5, but not in other TRPC members; 3) a pre-S1 elbow helix connects the N terminus and TMD in TRPC4 (**Fig. 2d**), as in TRPM4 and NOMPC (19, 22); however, TRPC4 and TRPM4’s pre-S1 helix is not found in NOMPC). In TRPC4 the pre-S1 elbow helix directly connects to the pre-S1 helix, while in TRPM4 a characteristic “bridge loop” (approximately 60 residues) connects the pre-S1 helix with the pre-S1 elbow (**Fig. 2d**).

**Figure 2:**
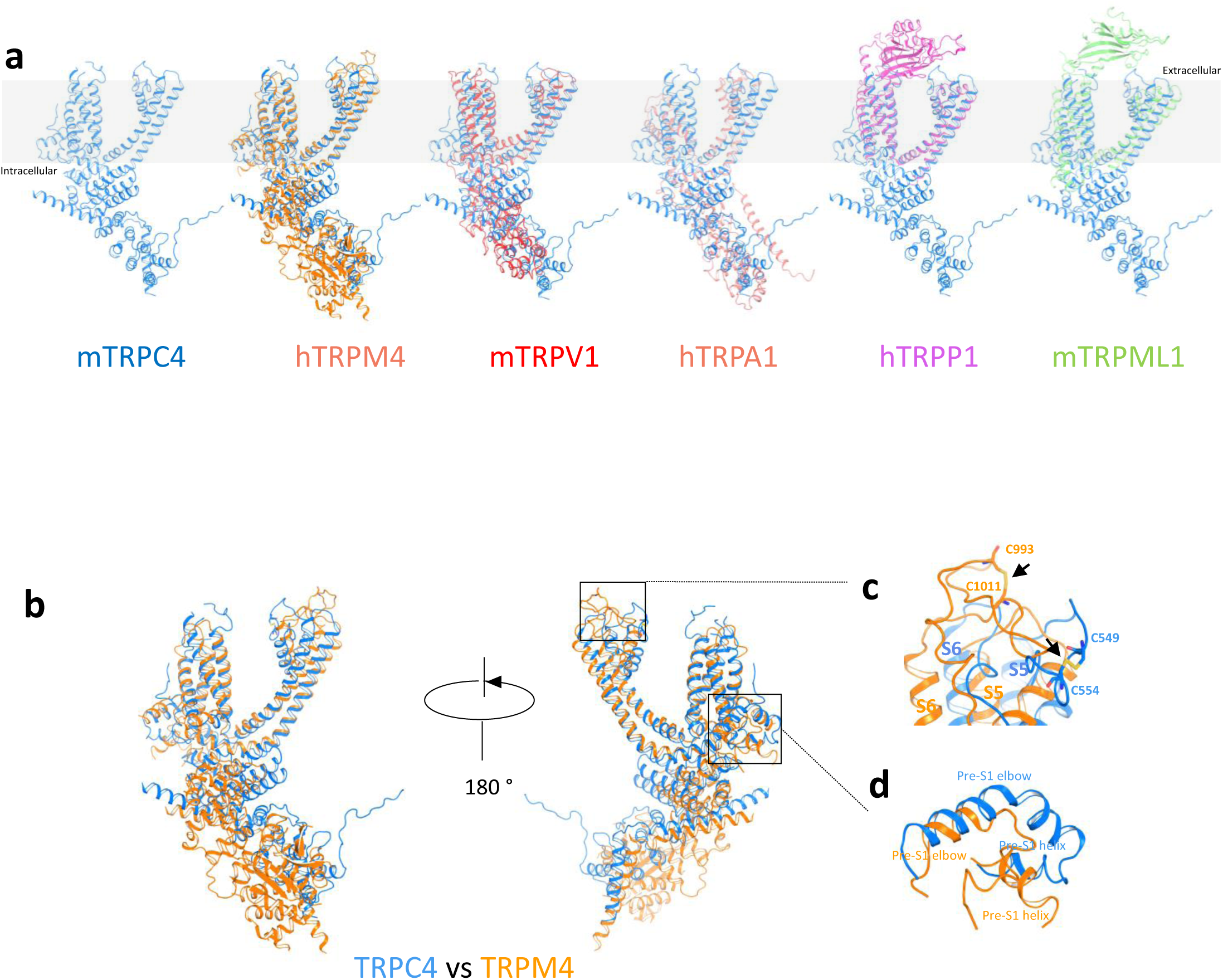
Comparison of the TRPC4 structure with previously solved TRP channel structures (apo state) **a**, Side views of an mTRPC4 subunit compared with other TRP family members including human TRPM4 (PDB: 6BWI)(20), mouse TRPV1 (PDB: 3J5P)(13), human TRPA1 (PDB: 3J9P)(16), human PKD2/TRPP1 (PDB: 5T4D)(17), and mouse TRPML1 (PDB: 5WPV)(18). **b**, Superimposition of TRPC4 and TRPM4. **c**, Key pore loop-disulfide bond between Cys549 and Cys554 in TRPC4 and the corresponding pore loop-disulfide bond between Cys993 and Cys1011 in TRPM4 (Black arrows); **d**, Differences in the organizations of the linker (pre-S1 elbow and pre-S1 helix) between the N terminus and transmembrane domains in TRPC4 and TRPM4.

### Cytosolic domain features and interactions

The cytosolic domains of TRP channels include regulatory components and domain interactions that may tune channel gating. The cytosolic domain of TRPC4 adopts a pedestal-like architecture (**Fig. 3a** and **Extended Data Fig. 5a**). The large and unique N-terminal domain of TRPC4 contains a long loop followed by an ankyrin repeat domain (ARD) and helix-loop-helix (HLH) motifs. These HLH motifs consist of seven helices and several connecting loops (**Fig. 1c, d** and **Extended Data Fig. 5a**). Similar to TRPM structures, the C-terminal domain of TRPC4 is composed of two helices, a connecting helix and a coiled-coil domain helix (**Fig. 1c, d**). The connecting and coiled-coil domain helices bend ∼120 degrees to form an inverted “L” architecture (**Extended Data Fig. 5b**). The coiled-coil domain contains three heptad repeats that exhibit the characteristic periodicity (a-b-c-d-e-f-g)_n_ (**Fig. 3 b, c** and **Extended Data Fig. 6**), with hydrophobic residues at positions “a” and “d”. The presence of Val and Ile at the “a” position, and Leu and Phe at the “d” position in the core of the coiled-coil domain supports the formation of a tetramer (**Fig. 3c**).

**Figure 3.**
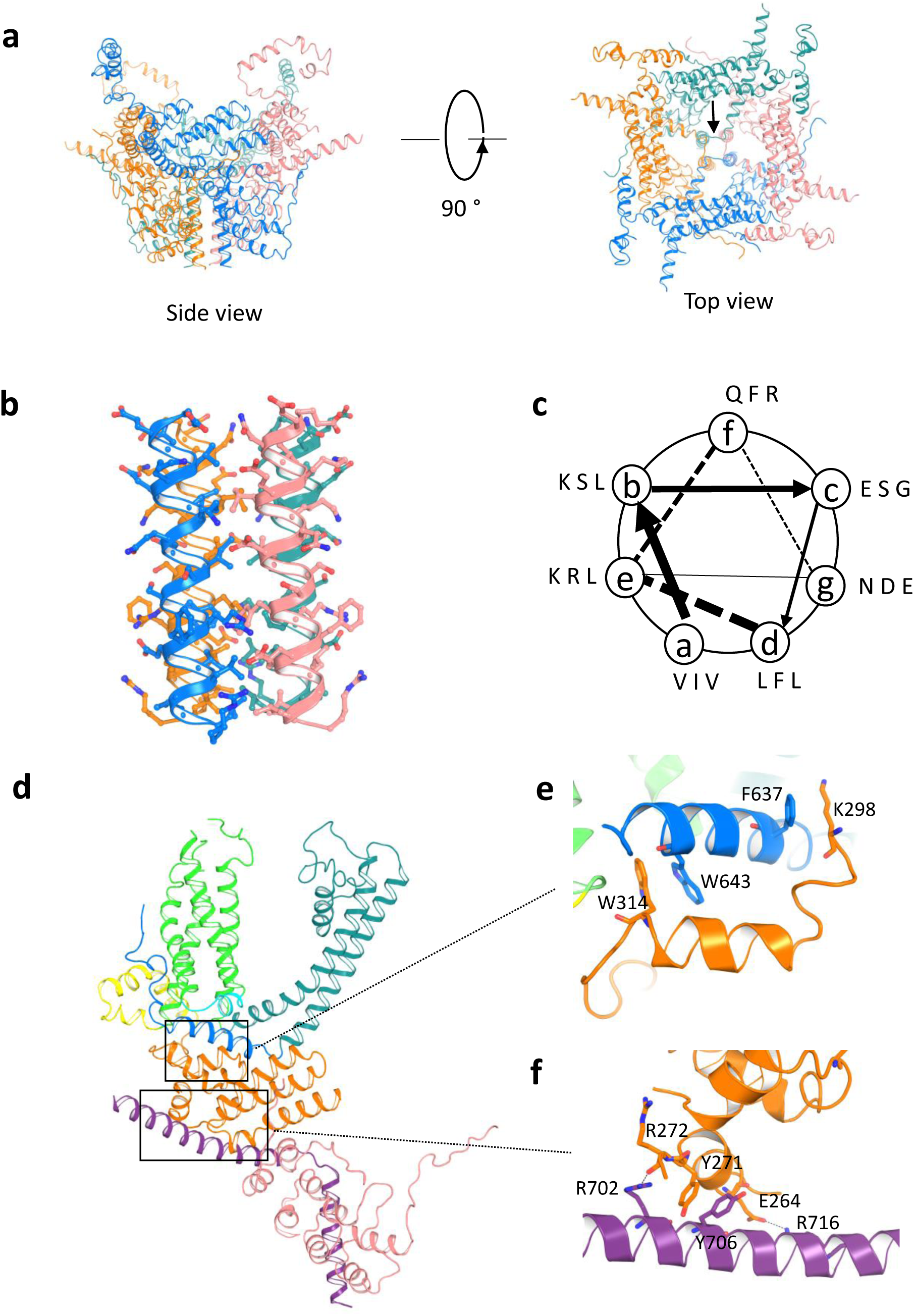
Unique cytosolic domains and interactions. **a**, Side and top views of the cytosolic domains including full-length N- and truncated C-terminal domains. **b**, Ribbon diagram of the tetrameric coiled-coil domain structure. Side chains are represented by ball and stick models. **c**, Helical wheel projection of the residues in the coiled-coil domain of TRPC4. **d**, Side views of a single subunit of the N-terminal domain to illustrate the locations of the interactions between the **e**, TRP domain (*Blue*) and N-terminal domain (*orange*) and **f**, N-terminal domain (*orange*) and truncated C-terminal domain (*purple*).

Aromatic interactions are important in cytosolic domain arrangements and protein folding (23). The TRP domain and N-terminal domain interactions are stabilized by π-π interactions (formed by Trp643 with Trp314) and cation-π interactions (formed by Phe637 with Lys298; **Fig. 3d, e**). The N- and C-terminal domains interface is also strengthened by a π-π interaction (Tyr271 with Tyr706) and two hydrogen bonds (Glu264 with Arg716, Arg272 with Arg702) (**Fig. 3d, f**).

### The ion conduction pore, cation and lipids binding sites

Positioned C-terminal to the pore helix, Gly577 marks a restriction point of 6.7 Å between diagonally opposed residues (Fig. 4 a, b). The corresponding filter-forming residue in TRPM4 is Gly976 at a 6.0 Å constriction. Compared to TRPM4, TRPC4’s selectivity filter is slightly more open, but the ion conduction pathway is restricted at its cytoplasmic interface, with Ile617, Asn621, and Gln625 at the bottom of S6 defining a lower gate. The narrowest constriction of the ion conduction pathway (3.6 Å) is formed by the S6 side chains of Asn621 (Fig. 4c and **Extended Data Fig. 7a**), while in TRPM4, the 5.1 Å wide lower gate is positioned at Ile1040 (**Extended Data Fig. 7b**). In contrast, the most restricted point in TRPV1 is in the selectivity filter (4.8 Å) between opposing Gly643 residues (**Extended Data Fig. 7c**) (24). In TRPA1, the narrowest point (6.1 Å) is Val961, which is found at its lower gate (16) (**Extended Data Fig. 7d**). These ∼0.5-2.5 Å differences in the narrowest point of TRPs structures may give some clue as to ion selectivity and activation mechanisms, but we also are aware that current resolution optimization in cryo-EM is still being improved by methods such as model-based local density sharpening (25), and resolution varies with location within the particle, conditions such as vitrification, and electron density map fitting.

**Figure 4.**
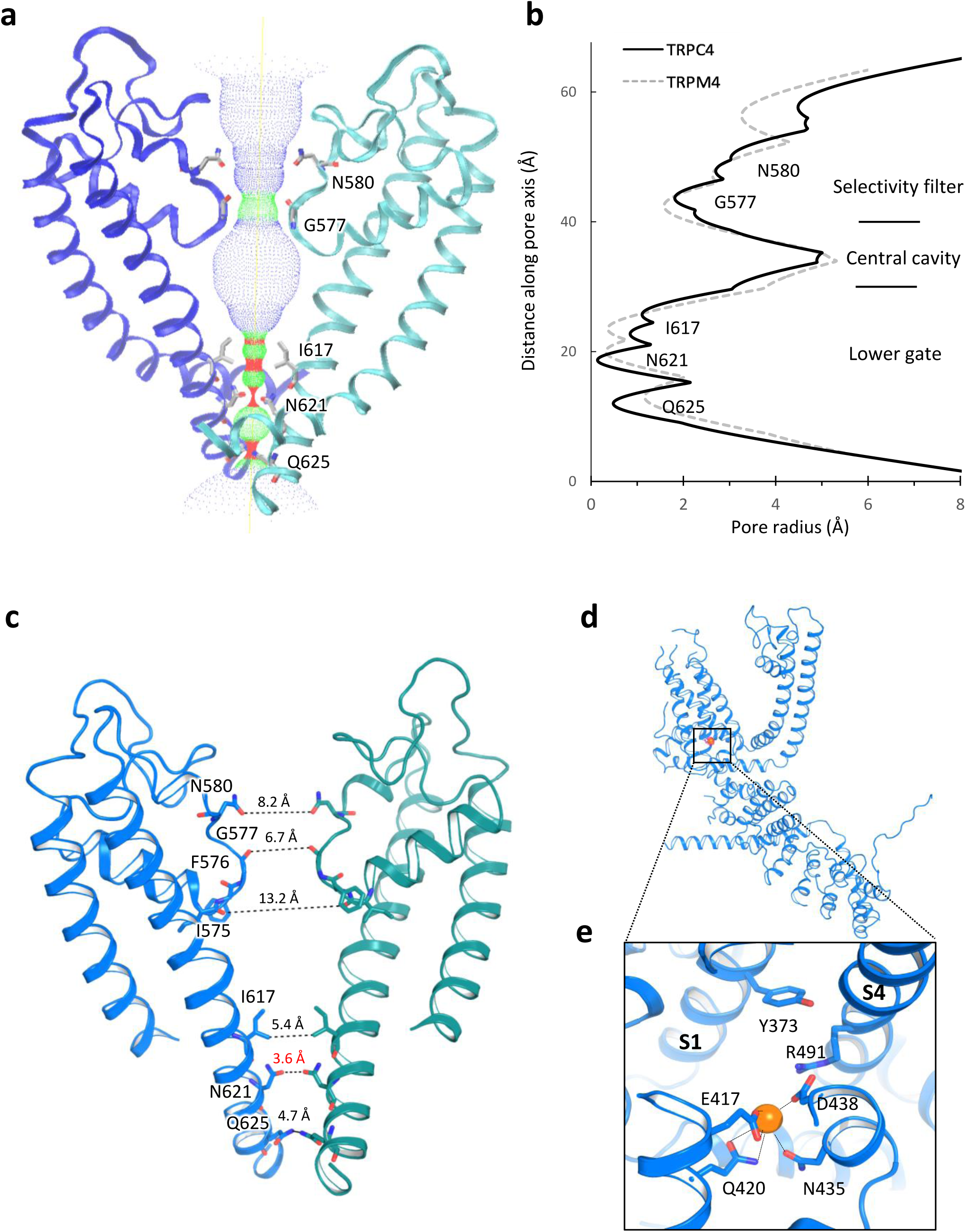
TRPC4 ion conduction pathway. **a**, Ion conduction pathway shown as dots and mapped using HOLE. **b**, Pore radius along the central axis. The side chains of G577 form a narrow constriction at the selectivity filter. N621 is the most restricted site of the lower gate. The dashed line represents TRPM4 for comparison. **c**, Side view of TRPC4’s pore region with chains A and C. The distances between diagonal residues in the selectivity filter and lower gate are labeled. **d**, A putative Na^+^ is found on the cytosolic side in the hydrophilic pocket of the S1-S4 domain interacting with E417, Q420, N435 and D438 (orange sphere). **e**, Enlarged view of second putative Na^+^ binding site.

The simplest hypothesis, with these caveats in mind, is that TRPC4 is in a closed or inactivated state since the lower gate is too narrow to allow the passage of a fully or partially hydrated ion. In support of this idea is the fact that Gln625 (located in the ion conduction exit pathway) is conserved in all the TRPC channels, suggesting it plays an important role in ion permeation (**Extended Data Fig. 8**).

TRPC4 is non-selective and thus permeable to monovalents (Na^+^, K^+^) and some divalents, such as Ca^2+^. A strong non-protein density peak in our TRPC4 structure is present in a hydrophilic pocket on the cytoplasmic side of the S1-S4 fold, consistent with the corresponding location of a presumed Ca^2+^ in TRPM4 (**Fig. 4d, e** and **Extended Data Fig. 9**) (19). We tentatively modelled this non-protein density as Na^+^ since sodium was the most abundant cation in our purification buffer. The assumed Na^+^ located at the cytoplasmic face is apparently coordinated by side chains of Glu417 and Gln420 from S2 and the Asp438 and Asn435 from S3 (**Fig. 4e**). The negatively charged Glu417 and Asp438 are conserved within the TRPC subfamily (except TRPC1) (**Extended Data Fig. 8**). S1’s Tyr373 and the positively charged S4 Arg491 are located above the cation binding site, forming a lid that may prevent the outward movement of cations (**Fig. 4e)**. Eight densities corresponding to lipid molecules were clearly resolved and identified as cholesteryl hemisuccinates (CHS) and phospholipids (the density fitting ceramide-1-phosphate or phosphatidic acid) (**Extended Data Fig. 3** and **Extended Data Fig. 10**). Four CHS located at the interface of the N-terminal domain and the S4/S5 linker are bound to each protomer, stabilizing the domain interaction (**Extended Data Fig. 10**). The phospholipid is embedded in the gap between the 4 monomeric subunits with its polar head interacting with the pore helix and neighboring S6 helix (**Extended Data Fig. 10**). In vivo phosphorylation or dephosphorylation of membrane lipids could thus alter the topology of the ion conduction pathway.

The TRPC4 structure provides a detailed view of the core domain of the canonical TRPC subfamily. Along with the other recent TRP structures, we now have general structural principles of this family of proteins. Comparison with other TRP channel structures highlights some commonalities and differences. Not surprisingly, all TRP channels are tetramers with domain swapping interactions, pore loops, selectivity filters, and extracellular and intracellular-facing constriction sites, as first shown for 6 TM K^+^ channels (26). One interesting feature that bears functional investigation is the extracellular pore loop disulfide bond (e.g., TRPC4 and TRPM4). Interestingly, the lower gate in the TRPC4 appears to have an unusual set of three constriction sites not found in other TRP channel structures. An ion binding site located in the hydrophilic binding pocket in the S1-S4 domain is observed in TRPC4 and TRPM4, which may be a general feature of these two subfamilies. We suspect that the most interesting differences between TRP channels lies in their less well structurally and functionally characterized extracellular and intracellular domains. These areas are best suited to ligand interactions that alter gating and drive the evolution of the ∼30 TRP channel members.

## Materials and Methods

### Protein expression and purification

The mouse TRPC4 construct (a.a. 1-758 of 974), was cloned into the pEG BacMam vector (27) and a maltose binding protein (MBP) tag was added to its N terminus. P3 baculovirus were produced in the Bac-to-Bac Baculovirus Expression System (Invitrogen). HEK293S GnTI-cells were infected with 10% (v/v) P3 baculovirus at a density of 2.0 − 3.0 × 10^6^ cells/ml for protein expression at 37°C. After 12-24 h, 10 mM sodium butyrate was added and the temperature reduced to 30°C. Cells were harvested at 72 h after transduction, and resuspended in a buffer containing 30 mM HEPES, 150 mM NaCl, 1 mM dithiothreitol (DTT), pH 7.5 with EDTA-free protease inhibitor cocktail (Roche). After 30 min, cells were solubilized for 2-3 h in a buffer containing 1.0 % (w/v) N-dodecyl-beta-D-maltopyranoside (DDM, Affymetrix), 0.1% (w/v) cholesteryl hemisuccinate (CHS, Sigma), 30 mM HEPES, 150 mM NaCl, 1 mM DTT; pH 7.5 with EDTA-free protease inhibitor cocktail (Roche). The supernatant was isolated by 100,000×g centrifugation for 60 min, followed by incubation in amylose resin (New England BioLabs) at 4°C overnight. The resin was washed with 20 column volumes of ‘washing buffer’ containing 25 mM HEPES, 150 mM NaCl, 0.1% (w/v) digitonin, 0.01% (w/v) CHS, 1 mM DTT; pH 7.5 with EDTA-free protease inhibitor cocktail (Roche). The protein was eluted with 4 column volumes of washing buffer with 40 mM maltose. The protein was then concentrated to 0.5 ml with a 100 kDa molecular weight cut-off concentrator (Millipore). PreScission protease was added to the samples and incubated overnight at 4°C to remove the MBP tag. After incubation at 4°C overnight, the protein was then purified on a Superose 6 column in a buffer composed of 25 mM HEPES, 150 mM NaCl, 0.1% (w/v) digitonin, 1 mM DTT; pH 7.5. The peak, corresponding to tetrameric TRPC4 was collected and concentrated to 4.5 mg/ml for cryo-EM study.

### Electron microscopy data collection

Purified TRPC4 protein (3.5 µl) in digitonin at 4.5 mg/ml was applied onto a glow-discharged, 400 mesh copper Quantifoil R1.2/1.3 holey carbon grid (Quantifoil). Grids were blotted for 7 s at 100% humidity and flash frozen by liquid nitrogen-cooled liquid ethane using a FEI Vitrobot Mark I (FEI). The grid was then loaded onto an FEI TF30 Polara electron microscope operated at 300 kV accelerating voltage. Image stacks were recorded on a Gatan K2 Summit (Gatan) direct detector set in super-resolution counting mode using SerialEM (28), with a defocus range between 1.5 to 3.0 μm. The electron dose was set to 8 e^-^/physical pixel/s and the sub-frame time to 200 ms. A total exposure time of 10 s resulted in 50 sub-frames per image stack. The total electron dose was 52.8 e^-^ per Å^2^ (∼1.1 e^-^ per Å^2^ per sub-frame).

### Image processing and 3D reconstruction

Image stacks were gain-normalized and binned by 2× to a pixel size of 1.23 Å prior to drift and local movement correction using motionCor2 (29). The images from the sum of all frames with dose-weighting were subjected to visual inspection and poor images were removed before particle picking. Particle picking and subsequent bad particle elimination through 2D classification was performed using Python scripts/programs (30) with minor modifications in the 8x binned images. The selected 2D class averages were used to build an initial model using the common lines approach implemented in SPIDER (31) through Maofu Liao’s Python scripts (30), which was applied to later 3D classification using RELION (32). Contrast transfer function (CTF) parameters were estimated using *CTFFIND4* (33) using the sum of all frames without dose-weighting. Quality particle images were then boxed out from the dose-weighted sum of all 50 frames and subjected to RELION 3D classification. RELION 3D refinements were then performed on selected classes for the final map. The resolution of this map was further improved by using the sum of sub-frames 1-14.

### Model building, refinement and validation

For the TRPC4, a polyalanine model was first built in COOT (34). Taking advantage of the defined geometry of helices and clear bumps for Cα atoms in the transmembrane domain, amino acid assignment was subsequently achieved based primarily on the clearly defined side chain densities of bulky residues. The refined atomic model was further visualized in COOT. A few residues with side chains moving out of the density during the refinement were fixed manually, followed by further refinement. The TRPC4 model was then subjected to global refinement and minimization in real space using the PHENIX (35) module ‘phenix.real_space_refine’(36) and geometry of the model was assessed using MolProbity (37) in the comprehensive model validation section of PHENIX. The final model exhibited good geometry as indicated by the Ramachandran plot (preferred region, 97.39%; allowed region, 2.08%; outliers, 0.53%). The pore radius was calculated using HOLE (38).

### Electrophysiology and Ca^2+^ measurements

TRPC4 constructs or empty vector were transfected into 293T cells together with an mCherry plasmid. Cells with red fluorescence were selected for whole-cell patch recordings (HEKA EPC10 USB amplifier, Patchmaster 2.90 software). A 1-s ramp protocol from –100 mV to +100 mV was applied at a frequency of 0.2 Hz. Signals were sampled at 10 kHz and filtered at 3 kHz. The pipette solution contained (mM): 140 CsCl, 1 MgCl_2_, 0.03 CaCl_2_, 0.05 EGTA, 10 HEPES, and the pH was titrated to 7.2 using CsOH. The standard bath solution contained (mM): 140 NaCl, 5 KCl, 1 MgCl_2_, 2 CaCl_2_, 10 HEPES, 10 D-Glucose, and the pH was adjusted to 7.4 with NaOH. The recording chamber had a volume of 150 µl and was perfused at a rate of ∼2 ml/min. For Ca^2+^ imaging experiments, transfected 293T cells were seeded on coverslips and incubated with Fura-2 AM (2 μM) for 30 min at 37°C in standard bath solution. The ratio (F_340_/F_380_) of Ca^2+^ dye fluorescence was measured by a Nikon Ti-E system with NIS-Elements software. All the experiments were performed at room temperature.

## Acknowledgements

We thank Dr. Steve Harrison and the Cryo-EM Facility (Harvard Medical School) for use of their microscopes. We thank Dr. Maofu Liao for providing the Python scripts and help in image processing. We thank Dr. Corey Valinsky’s help on manuscript revision. J.Z. was supported by the Thousand Young Talents Program of China and National Natural Science Foundation of China (Grant No. 31770795). J.L.was supported by the National Natural Science Foundation of China (Grant No. 81402850). Functional studies in this project were supported by the National Natural Science Foundation of China (31300949 to B.Z. and 31300965 to G.L.C.)

## Author Contributions

J.Z. and J.D. designed and made constructs for BacMam expression and determinded the condition to enhance protein stability. J.Z. purified the protein. Z.L. carried out detailed cryo-EM experiments, including data acquisition and processing. J.L., and J.Z. built the atomic model on the basis of cryo-EM maps. B.Z. and G.L.C. performed functional studies. X. P., Y.Z. and J.W. assisted with protein purification and the mutation of TRPC4 constructs for functional studies. J.D. and J.Z. drafted the initial manuscript. All authors contributed to structure analysis/interpretation and manuscript revision. J.Z. and Z.L. initiated the project, planned and analyzed experiments and supervised the research.

The authors declare no competing financial interests.

### Data deposition

Cryo-EM electron density map of the mouse TRPC4 has been deposited in the Electron Microscopy Data Bank, https://www.ebi.ac.uk/pdbe/emdb/ (accession number EMD-6901), and the fitted coordinate has been deposited in the Protein Data Bank, www.pdb.org (PDB ID code 5Z96).

## Extended Data Figure

**Extended Data Figure 1. The TRPC4 construct encodes a functional channel; biochemical characterization.**

**a**, Size exclusion chromatography trace of TRPC4 proteins. Void volume (V_0_) and the peaks corresponding to tetrameric TRPC4 and MBP are indicated. Protein samples of the indicated TRPC4 protein fraction were subjected to SDS-PAGE and Coomassie-blue staining. **b**, Intracellular Ca^2+^ measurements as indicated by Fura-2 AM (2 μM). **c**, Representative whole-cell patch clamp recordings and *I-V* relationships of truncated mTRPC4, full-length mTRPC4, and empty vector expressed in HEK293T cells. TRPC4 sensitivity to the activator Englerin A, blockers ML204 and 2-APB, and the GPCR agonist trypsin was not affected by truncation.

**Extended Data Figure 2. Flow chart for cryo-EM data processing of the TRPC4 structure. a**, Representative image of the purified TRPC4 protein, 2D class averages of TRPC4 particles, side views of the 3D reconstructions from RELION 3D classification and final 3D reconstructions from 3D auto-refinement. **b**, Fourier shell correlation (FSC) curve for the 3D reconstruction (marked at overall 3.3 Å resolution). **c**, Local resolution estimation from ResMap (39) and **d**, Euler distribution plot of particles used in the final three-dimensional reconstruction. The length of the rod is propotional to the number of particles in that view, with regions in red denoting the views containing the highest number of particles.

**Extended Data Figure 3. Cryo-EM densities of selected regions of TRPC4.**

Density map showing the transmembrane helices (S1-S6), an ankyrin repeat (AR), N-terminal helix, TRP domain, pore helix, connecting helix, coiled-coil helix, and lipids. The maps were contoured at a level of 3.0 σ.

**Extended Data Figure 4. Comparison of the 6 transmembrane domain structures of TRPC4 and TRPM4.**

Side (left) and top (right) views of the channel transmembrane domain monomers of two channels were overlapped for comparison. The helices of the apo states of TRPC4 (blue) and TRPM4 (orange) adopt a similar conformation in S1-S4, but differ in the S2-S3 linker and the orientation of S5 and S6.

**Extended Data Figure 5. Cytosolic domains of the TRPC4 monomer.**

Side views of the **a**, N-terminal and **b**, truncated C-terminal domains.

**Extended Data Figure 6. The three heptad repeats of the coiled-coil domain.**

**a**, Side and top views of the periodic region of the coiled-coil domain denoted as (a-b-c-d-e-f-g)_n_; **b**, Protein sequences of conserved coiled-coil domain of TRPC4 and TRPC5. Residues included in TRPC4 and TRPC5 are indicated in black. Residues at positions “a” and “d” are shown in red.

**Extended Data Figure 7. Comparison of ion conducting pathway in TRP family.**

Comparison of ion conduction pathway openings of **a**, TRPC4, **b**, TRPM4 (PDB: 6BWI), **c**, TRPV1 (PDB: 3J5P) and **d**, TRPA1 (PDB: 3J9P). Distances between specific side chains along the pore and the key residues are labeled.

**Extended Data Figure 8. Sequence alignment of TRPC subfamily members.**

Sequence of the full-length mouse TRPC4 aligned to other TRPC subfamily members are shown; key residues indicated. Regions corresponding to putative Na^+^ binding sites are labeled. The selectivity filter, lower gate, and two cysteines forming disulfide bonds are highlighted. Sequence alignments of this study were performed using Clustal Omega.

**Extended Data Figure 9. Electrostatic maps of the predicted Na**^**+**^ **binding sites.**

Side and top views of electrostatic maps of predicted Na^+^ binding pockets in TRPC4; **a**, monomer and **b**, tetramer. The surface is colored according to the calculated electrostatic potential. The electrostatics reveal the tetrameric distribution of charge. Blue indicates positive potential, red negative, and transparent white neutral.

**Extended Data Figure 10. Lipid coordination in TRPC4.**

**a**, Side and top views of ribbon diagrams of the TRPC4 tetramer: 4 cholesterol hemisuccinate (CHS) molecules and 4 phospholipids (potentially ceramide-1-phosphate, C1P, or phosphatidic acid, PA) shown in cyan. **b**, Side views of each CHS and PA molecules per protomer. **c** and **d**, Ribbon diagram of the TRPC4 lipids binding. **c**, CHS, shown in cyan, interacts with the S4/S5 linker and Tyr315 in the N-terminal domain. **d**, PA is imbedded in the gap between the pore helix and neighboring subunit and interacts with the head groups of Gln569, Trp573, and Ala598. **e**. Side view of the electrostatic map around the putative PA binding pocket. The surface is colored according to the calculated electrostatic potential, revealing the tetrameric distribution of charge. Blue shows positive potential, red negative, and transparent white neutral.

