## Supplementary Materials for "Structure of the mouse TRPC4 ion channel"

### ED Figure 1.

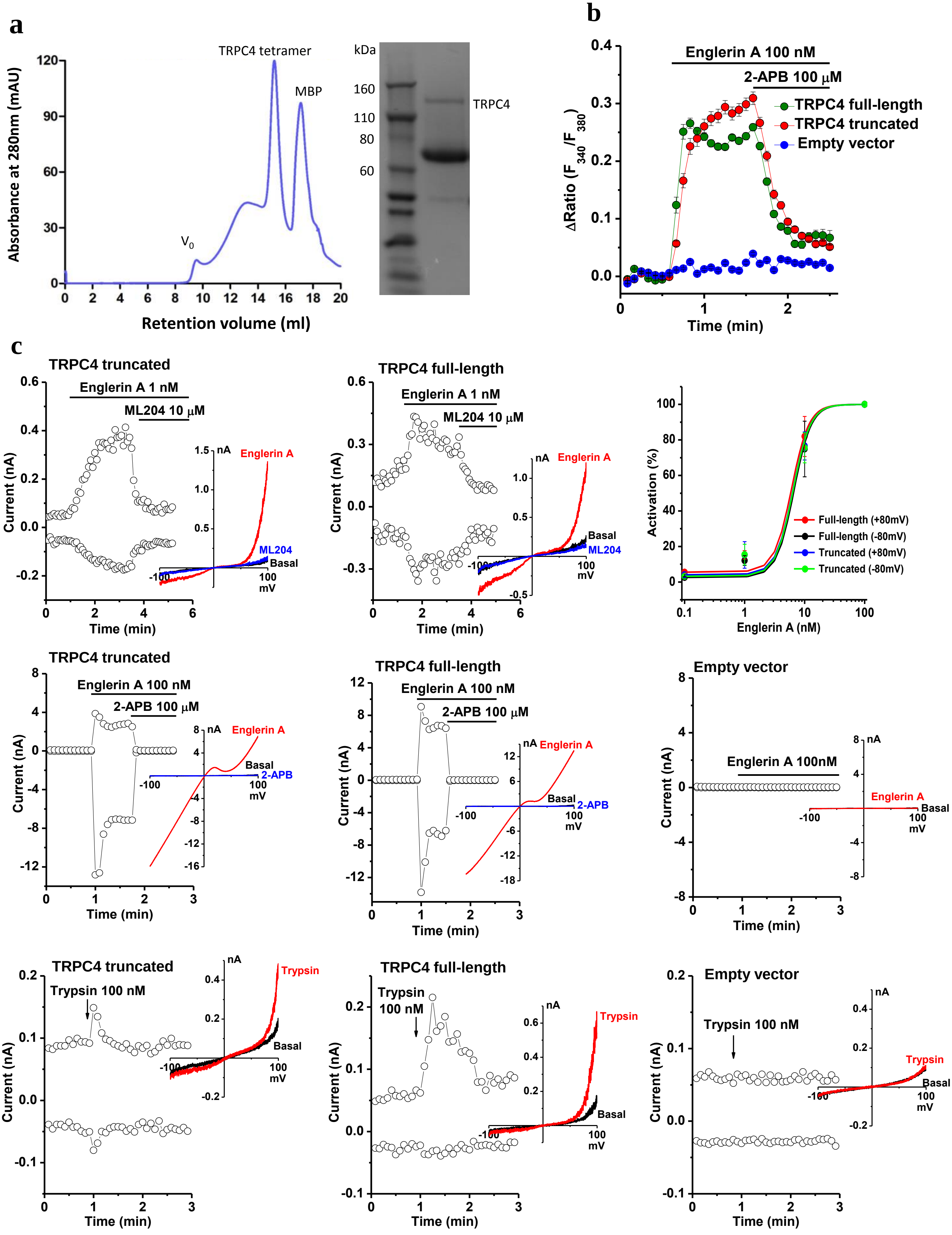

a
TRPC4 tetramer
MBP
V0
Absorbance at 280nm (mAU)
Retention volume (ml)
kDa
160
110
80
60
TRPC4
b
Englerin A 100 nM
2-APB 100 μM
TRPC4 full-length
TRPC4 truncated
Empty vector
ΔRatio (F340/F380)
Time (min)
c
TRPC4 truncated
Englerin A 1 nM
ML204 10 μM
Current (nA)
Time (min)
Englerin A
ML204
Basal
nA
mV
TRPC4 full-length
Activation (%)
Englerin A (nM)
Full-length (+80mV)
Full-length (-80mV)
Truncated (+80mV)
Truncated (-80mV)
Englerin A 100 nM
2-APB 100 μM
2-APB
Empty vector
Englerin A 100nM
Trypsin 100 nM
Trypsin

ED Figure 2.

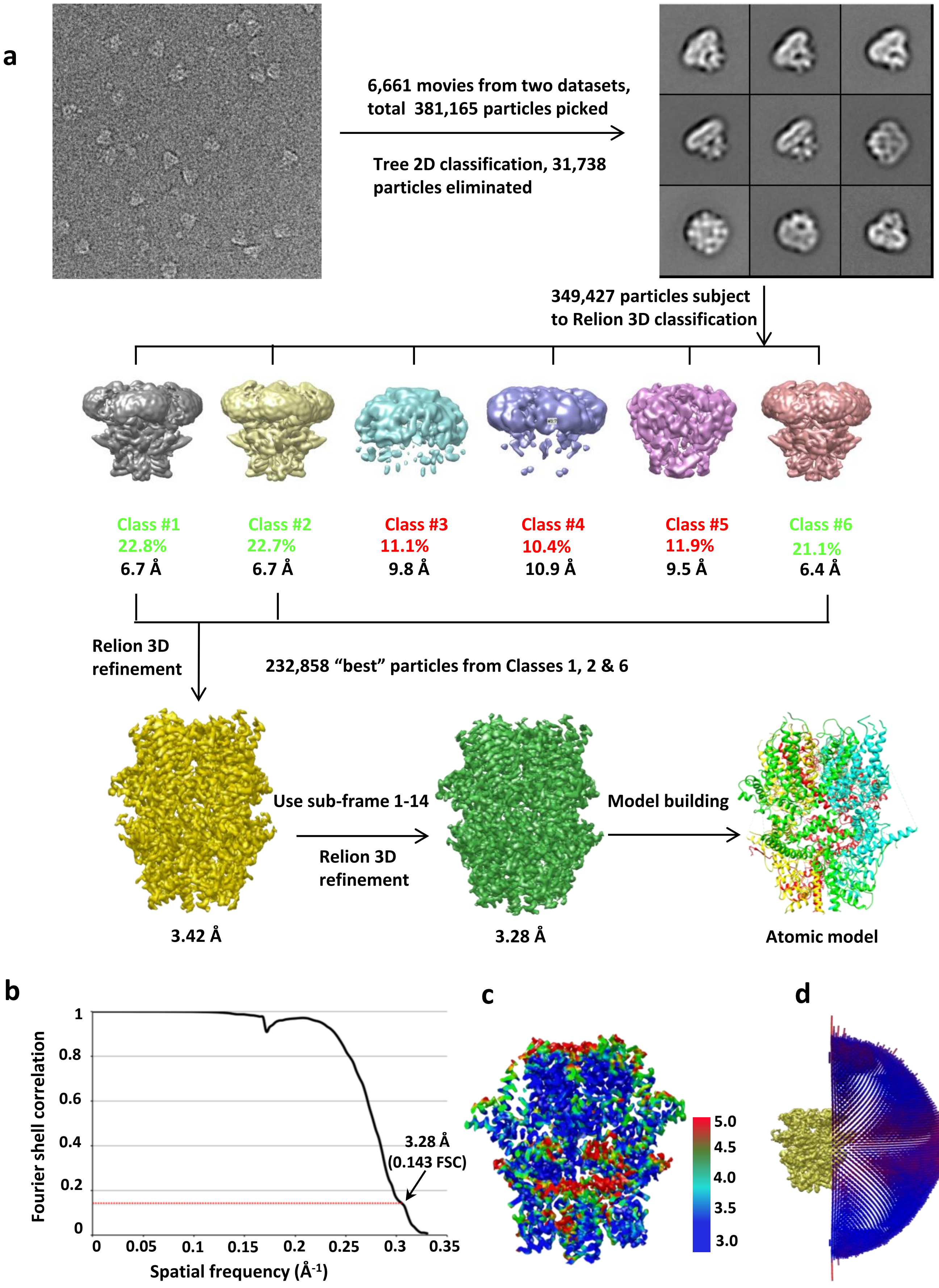
a
6,661 movies from two datasets, total 381,165 particles picked
Tree 2D classification, 31,738 particles eliminated
349,427 particles subject to Relion 3D classification
Class #1
22.8%
6.7 Å
Class #2
22.7%
6.7 Å
Class #3
11.1%
9.8 Å
Class #4
10.4%
10.9 Å
Class #5
11.9%
9.5 Å
Class #6
21.1%
6.4 Å
Relion 3D refinement
232,858 "best" particles from Classes 1, 2 & 6
Use sub-frame 1-14
Relion 3D refinement
Model building
3.42 Å
3.28 Å
Atomic model
b
Fourier shell correlation
Spatial frequency (Å-1)
1
0.8
0.6
0.4
0.2
0
0
0.05
0.1
0.15
0.2
0.25
0.3
0.35
3.28 Å
(0.143 FSC)
c
5.0
4.5
4.0
3.5
3.0
d

**ED Figure 3.**

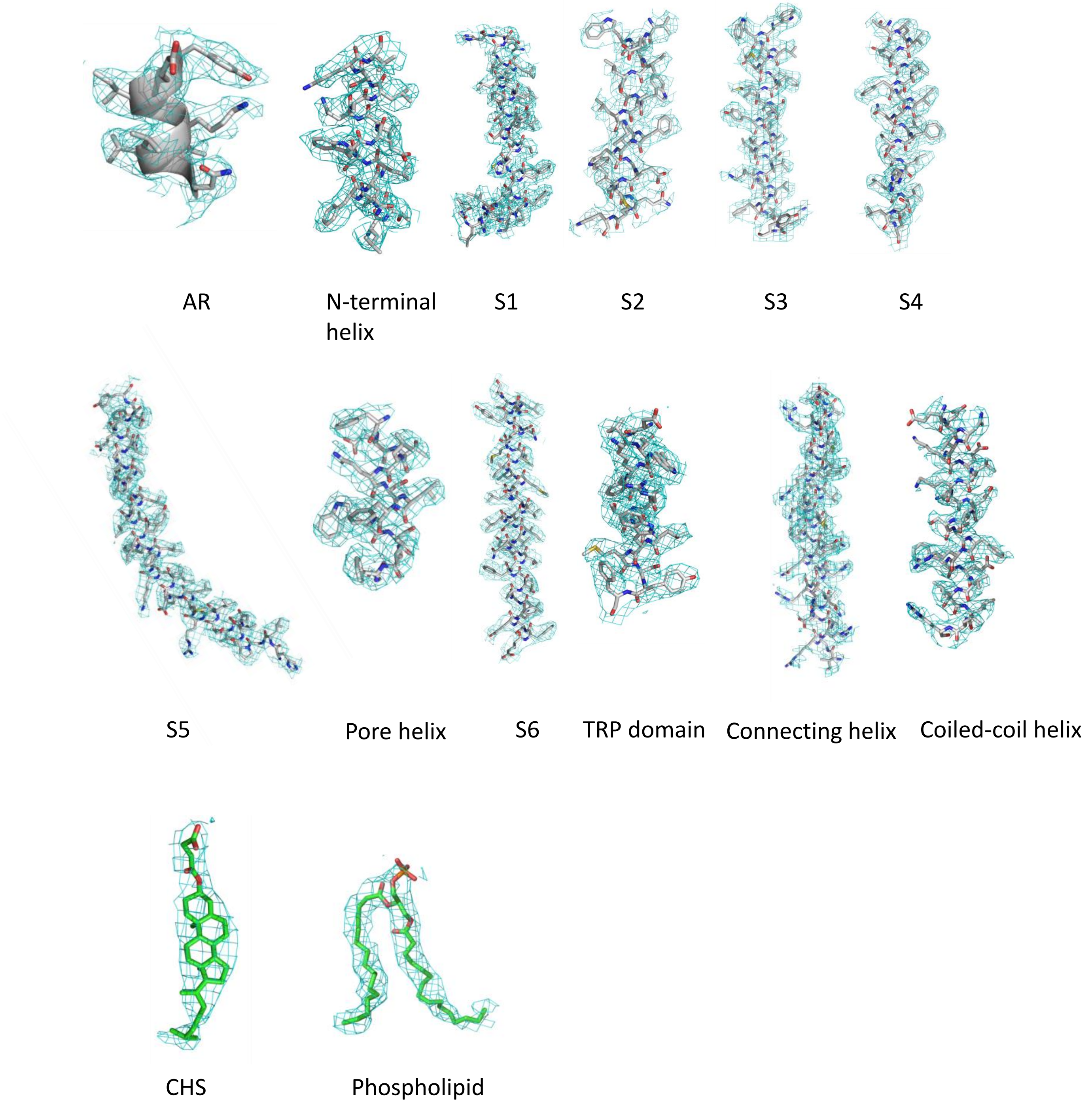
AR
N-terminal helix
S1
S2
S3
S4
S5
Pore helix
S6
TRP domain
Connecting helix
Coiled-coil helix
CHS
Phospholipid

### ED Figure 4.

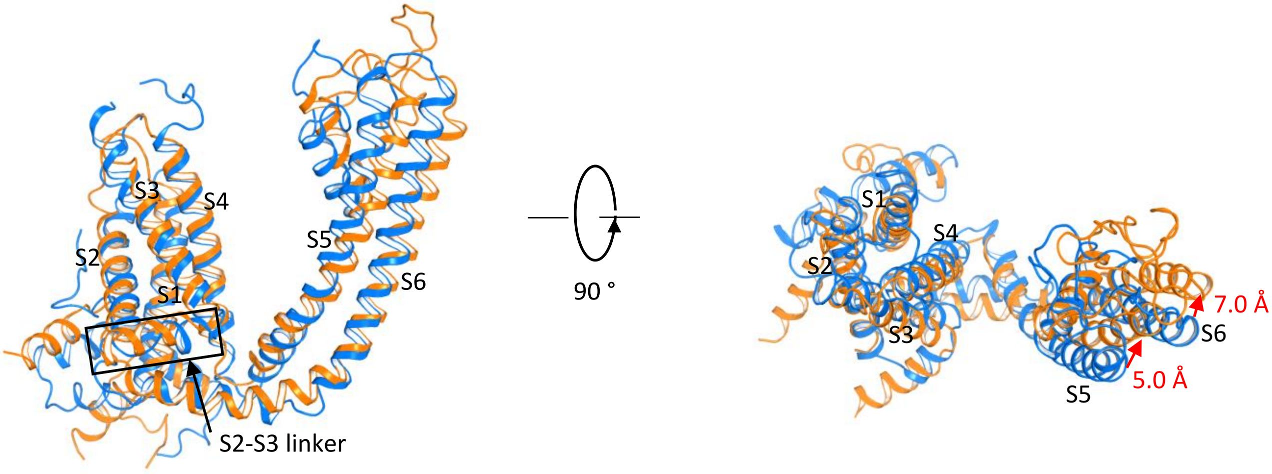

### ED Figure 5.

**a**

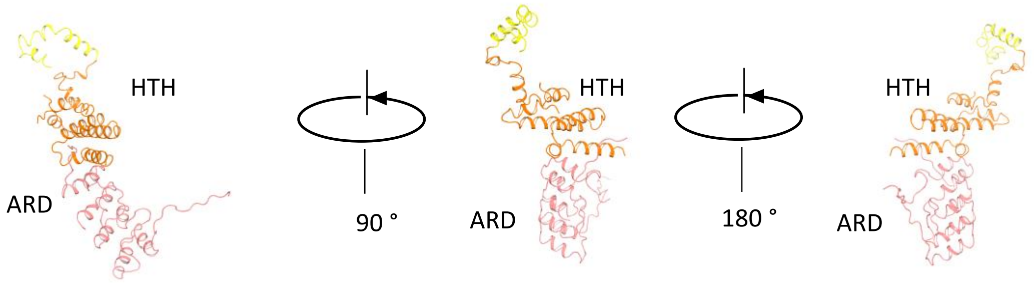


**b**

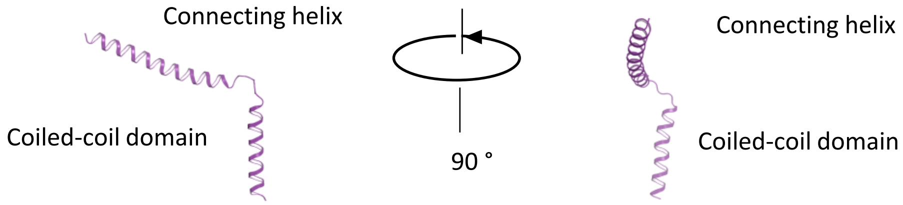

**ED Figure 6.**

**a**

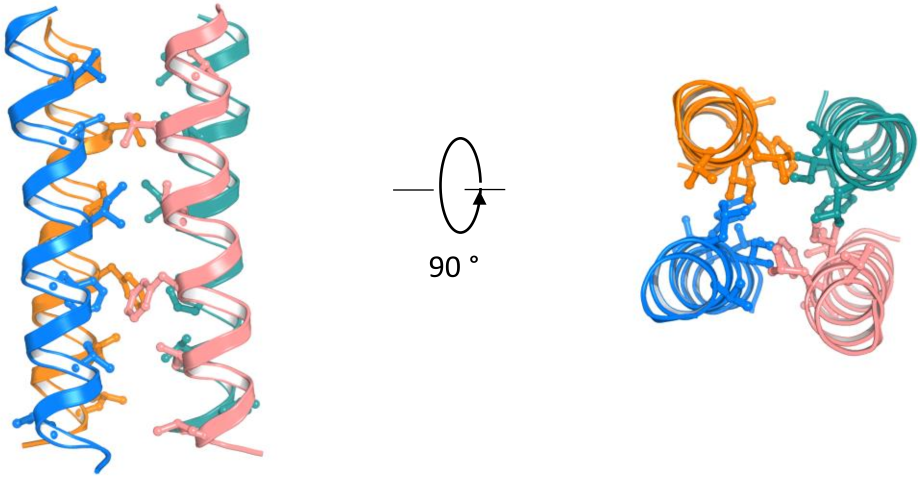


**b**

735 755

g a b c d e f g a b c d e f g a b c d e f

TRPC4 (mouse): T E E N V K E L K Q D I S S F R F E V L G L L R

TRPC5 (mouse): T E E N F K E L K Q D I S S F R Y E V L D L L G

**ED Figure 7.**

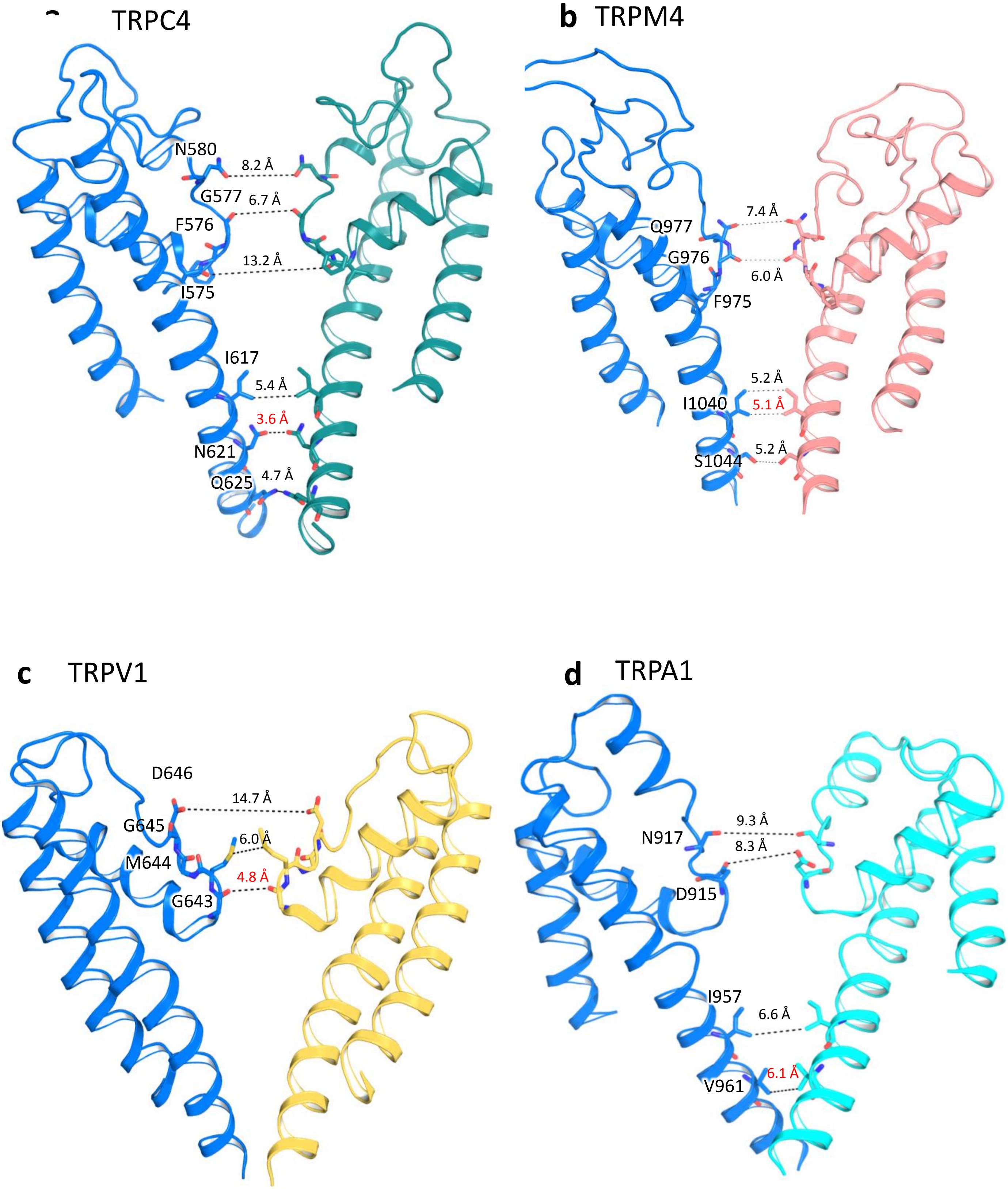

TRPC4
N580
8.2 Å
G577
6.7 Å
F576
13.2 Å
I575
I617
5.4 Å
3.6 Å
N621
Q625
4.7 Å
b TRPM4
7.4 Å
Q977
G976
6.0 Å
F975
5.2 Å
I1040
5.1 Å
S1044
5.2 Å
c TRPV1
D646
14.7 Å
G645
6.0 Å
M644
4.8 Å
G643
d TRPA1
9.3 Å
N917
8.3 Å
D915
I957
6.6 Å
6.1 Å
V961

### ED Figure 8.

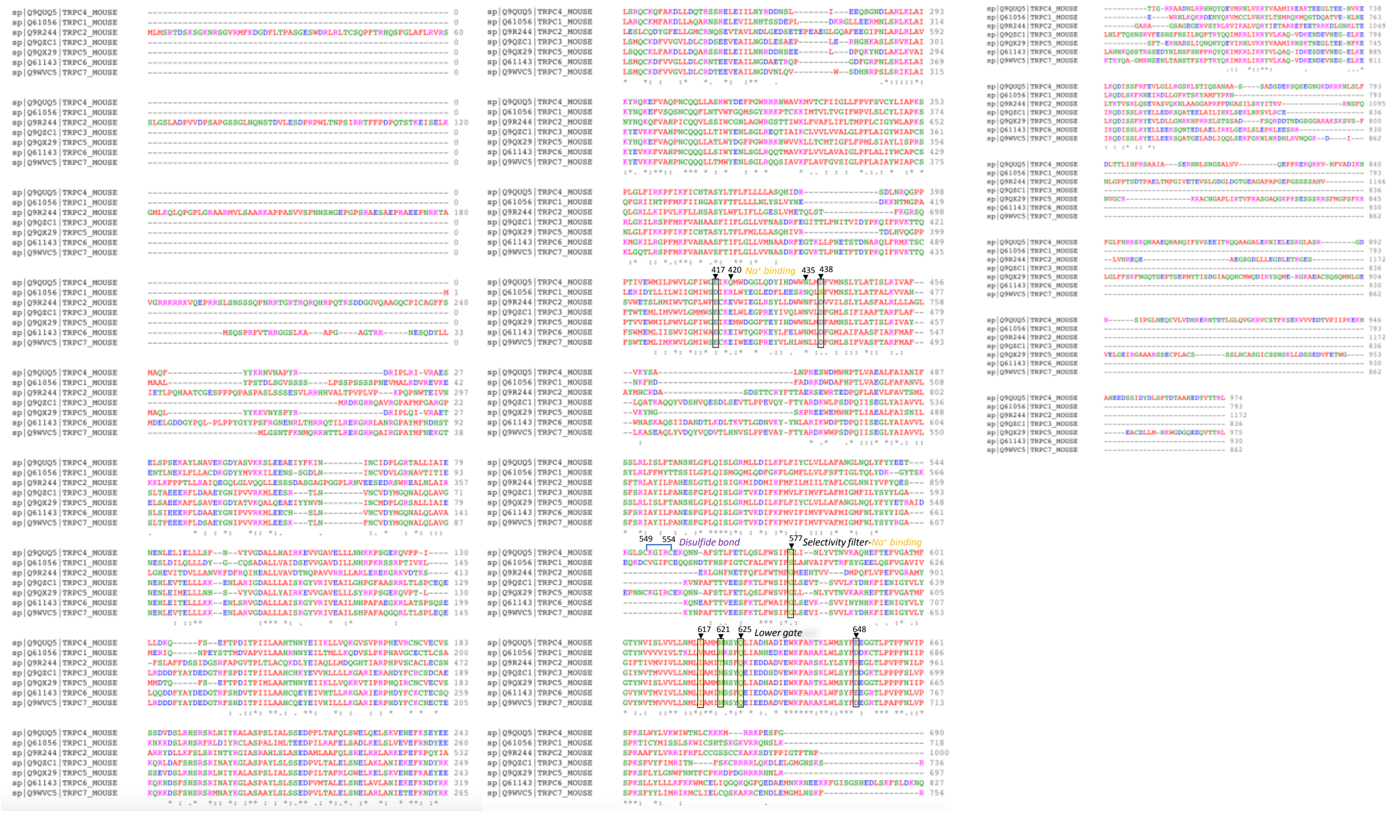
417 420
Na⁺ binding
435 438
549 554
Disulfide bond
577
Selectivity filter-Na⁺ binding
617 621 625
Lower gate
648

**ED Figure 9.**

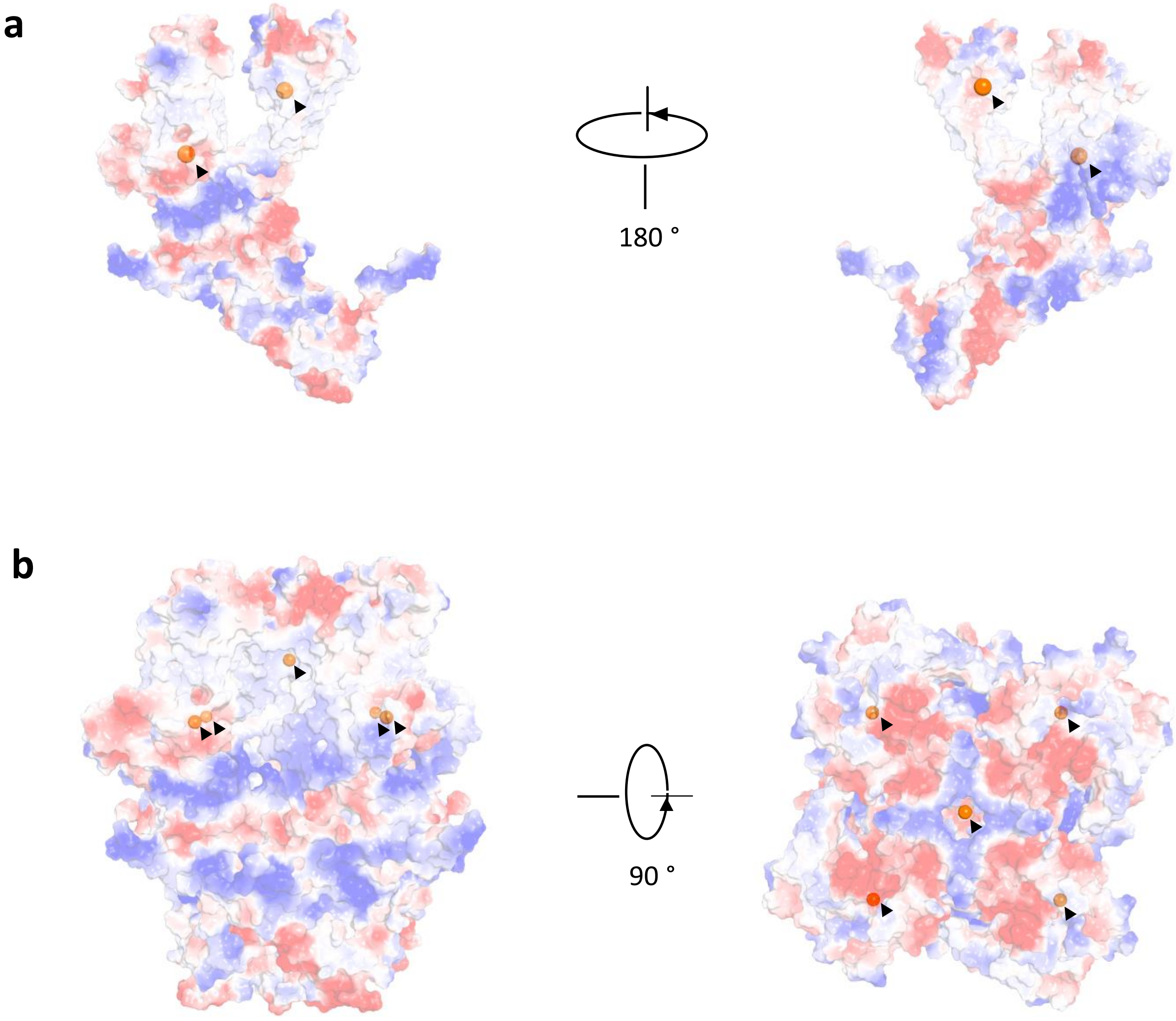

**ED Figure 10.**

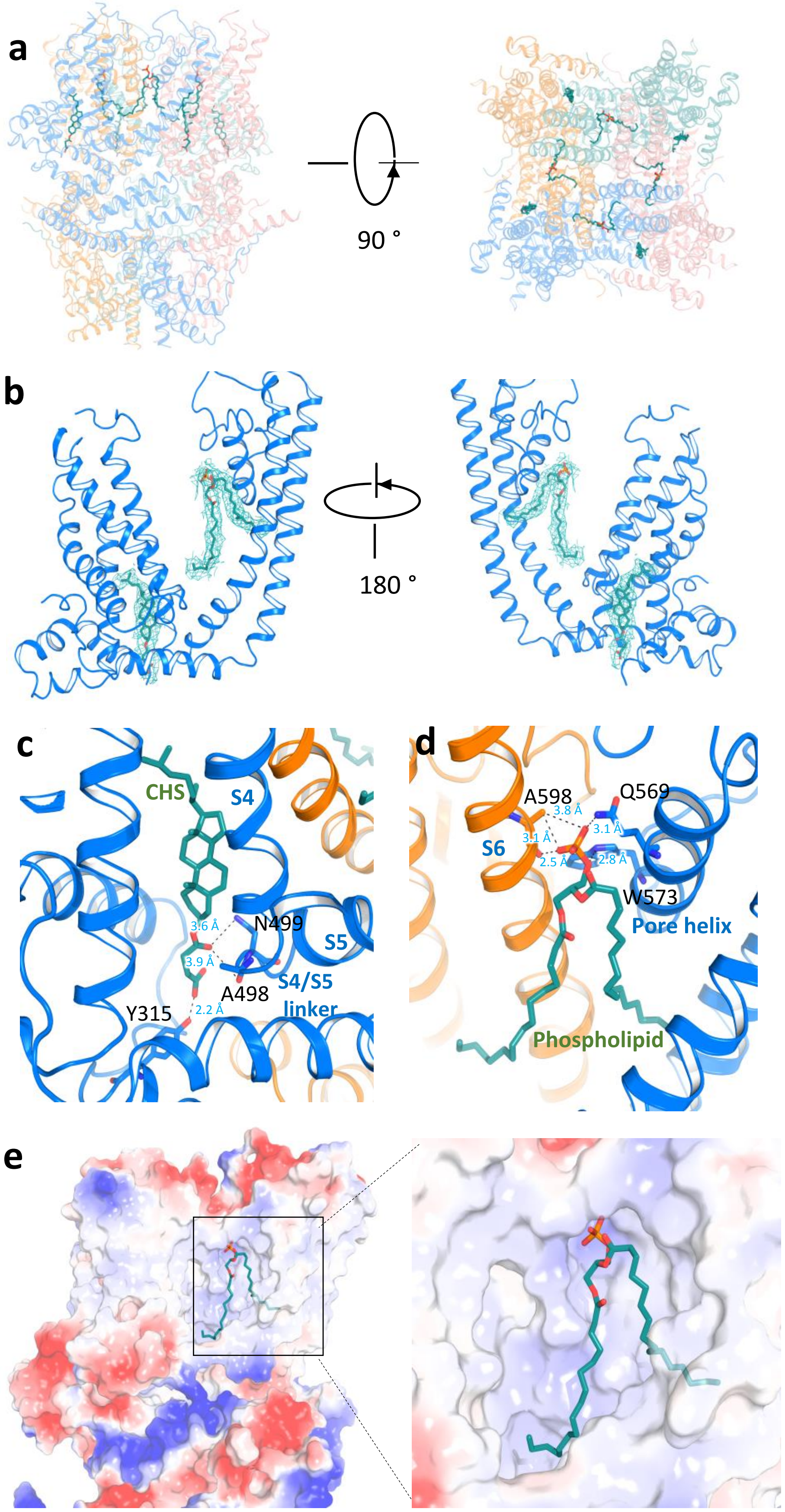
a
90 °
b
180 °
c
CHS
S4
S5
N499
A498
Y315
S4/S5
linker
3.6 Å
3.9 Å
2.2 Å
d
A598
Q569
S6
W573
Pore helix
Phospholipid
3.8 Å
3.1 Å
3.1 Å
2.5 Å
2.8 Å
e
