## Supplementary Materials for "Structure of the mouse TRPC4 ion channel"

**Table S1.** Cryo-EM data collection, refinement and validation statistics

| | MouseTRPC4 (EMD-6901) (PDB 5Z96) |
|---|---|
| **Data collection and processing** | |
| Magnification | 40,607 |
| Voltage (kV) | 300 |
| Electron exposure (e–/Å$^2$) | 52.8 |
| Defocus range (μm) | 1.5-3.0 |
| Pixel size (Å) | 1.23 |
| Symmetry imposed | *C4* |
| Initial particle images (no.) | 381,165 |
| Final particle images (no.) | 232,858 |
| Map resolution (Å) | 3.3 |
| **Refinement** | |
| Initial model used (PDB code) | *de novo* |
| Model resolution (Å) | 3.3 |
| Model composition | |
| Non-hydrogen atoms | 21,506 |
| Protein residues | 2,608 |
| Ligands | 13 |
| R.m.s. deviations | |
| Bond lengths (Å) | 0.01 |
| Bond angles (°) | 1.11 |
| Validation | |
| MolProbity score | 1.83 |
| Ramachandran plot | |
| Favored (%) | 97.39 |
| Allowed (%) | 2.08 |
| Disallowed (%) | 0.53 |
